# Upregulated miR-6818-5p in PCOS Patients Promotes Granulosa Cell Apoptosis and Suppresses Proliferation by Directly Targeting HSD17B2

**DOI:** 10.64898/2026.05.22.726113

**Authors:** Hai-Tao Pan, Feng Zhang, Hai-Gang Ding, Na Ding, Guo-Ping Li, Jin-Long Ding, Yao He, Tao Zhang, Xin-Yue Zhang, Bin Yu, Hong-Mei Lin

**Affiliations:** Shaoxing Maternity and Child Health Care Hospital, Shaoxing, China; Maternity and Child Health Care Affiliated Hospital, Shaoxing University; Shaoxing People’s Hospital (Shaoxing Hospital of Zhejiang University), Shaoxing, China

**Author notes:** **Corresponding Author:** Bin Yu, Shaoxing Maternity and Child Health Care Hospital, Shaoxing, China. E-mail address Or to Hong-Mei Lin, Shaoxing Maternity and Child Health Care Hospital, Shaoxing, China.

**Keywords:** Polycystic ovary syndrome, miR-6818-5p, HSD17B2, Granulosa cells, Apoptosis

## Abstract

Polycystic ovary syndrome (PCOS) is a prevalent endocrine disorder characterized by hyperandrogenism, ovulatory dysfunction, and polycystic ovaries, with granulosa cell dysfunction being a key pathological feature. This study aimed to investigate the role of microRNA-6818-5p in PCOS pathogenesis. Quantitative PCR revealed a significant upregulation of circulating miR-6818-5p in PCOS patients compared to healthy controls. In vitro, functional assays in the human granulosa cell line KGN demonstrated that miR-6818-5p overexpression markedly inhibited cell proliferation (assessed by CCK-8 assay) and promoted apoptosis (measured by Annexin V/PI flow cytometry). Mechanistically, dual-luciferase reporter assay and Western blotting identified HSD17B2 as a direct target of miR-6818-5p, with miR-6818-5p mimics significantly suppressing HSD17B2 protein expression. In conclusion, our findings reveal that elevated miR-6818-5p in PCOS may contribute to follicular development dysfunction by targeting HSD17B2 to disrupt granulosa cell proliferation and apoptosis balance, offering novel insights into PCOS pathology and highlighting miR-6818-5p as a potential diagnostic biomarker and therapeutic target.

## 1. Introduction

Polycystic ovary syndrome (PCOS) represents the most prevalent endocrine and metabolic disorder among women of reproductive age, with a global prevalence estimated between 5% and 11% [1]. This complex condition is clinically characterized by a constellation of features, including hyperandrogenism, ovulatory dysfunction, and polycystic ovarian morphology, which collectively contribute to significant reproductive morbidity, metabolic disturbances, and a diminished quality of life for affected individuals. The underlying pathophysiology of PCOS is multifaceted, involving intricate interactions between genetic predisposition, hormonal dysregulation, and environmental factors. Central to its reproductive pathology is the dysfunction of ovarian folliculogenesis, wherein the granulosa cells (GCs)—the somatic cells crucial for supporting oocyte development and steroidogenesis—exhibit aberrant proliferation, apoptosis, and hormonal responsiveness. This GC dysfunction is considered a pivotal contributor to the arrested follicular development and anovulation that hallmark the syndrome, yet the precise molecular drivers remain incompletely delineated[2-4].

In recent years, the role of post-transcriptional gene regulation, particularly mediated by microRNAs (miRNAs), has emerged as a critical layer of control in various physiological and pathological processes. miRNAs are small, non-coding RNA molecules that typically bind to the 3’-untranslated regions (3’-UTRs) of target messenger RNAs (mRNAs), leading to translational repression or mRNA degradation. Their involvement extends to key cellular functions such as proliferation, apoptosis, differentiation, and steroid hormone synthesis, all of which are relevant to ovarian physiology and PCOS pathogenesis. Notably, miRNAs are remarkably stable in bodily fluids, including blood, where they are protected within extracellular vesicles like exosomes, presenting a promising avenue for non-invasive biomarker discovery [1]. Consequently, there has been growing interest in profiling circulating miRNAs to identify novel diagnostic signatures for PCOS. For instance, sequencing analyses of serum exosomes have revealed distinct miRNA expression profiles in women with PCOS compared to healthy controls, highlighting their potential clinical utility [1].

Despite these advances, significant gaps persist in our understanding. The existing literature has identified several dysregulated miRNAs in the serum or follicular fluid of PCOS patients, such as miR-93, miR-132, and miR-145-5p, some of which have been linked to steroidogenic pathways or insulin signaling [1]. However, the functional consequences of most altered miRNAs, particularly their specific target genes and downstream effects on ovarian cell behavior, are not well characterized. Many studies remain correlative, lacking mechanistic validation in relevant cellular models. Furthermore, the PCOS miRNA landscape is likely far from fully mapped, with many miRNAs whose expression and function in the context of PCOS are entirely unknown[5-7]. This represents a critical knowledge gap, as elucidating the functional roles of specific miRNAs could uncover novel regulatory nodes in the pathological network of PCOS, bridging the gap between observed molecular alterations and the cellular dysfunction driving the disease.

To address this gap, the present study focuses on a specific miRNA, miR-6818-5p. Preliminary high-throughput sequencing data has suggested that miR-6818-5p is among the miRNAs differentially expressed in the serum exosomes of PCOS patients [1], but its expression pattern, clinical relevance, and biological function in PCOS have never been investigated. This miRNA represents a compelling candidate for several reasons. First, its identification in a sequencing profile of PCOS serum exosomes provides a direct, albeit preliminary, clinical link [1]. Second, while its roles in other contexts, such as cancer, hint at potential involvement in cell cycle regulation, its function in reproductive endocrinology is a complete mystery. Therefore, systematically investigating miR-6818-5p offers the opportunity to discover a previously unrecognized player in PCOS pathogenesis. The central hypothesis guiding this research is that miR-6818-5p is aberrantly expressed in PCOS and contributes to granulosa cell dysfunction by targeting key genes involved in cell survival and steroid metabolism.

To rigorously test this hypothesis, we designed an integrative study employing both clinical and cellular experimental approaches. Initially, we quantified the expression levels of miR-6818-5p in peripheral blood samples from a well-phenotyped cohort of PCOS patients and age-matched healthy controls using reverse transcription-quantitative polymerase chain reaction (RT-qPCR). To dissect its biological function, we utilized the human ovarian granulosa cell line, KGN, a validated model that retains many functional characteristics of primary granulosa cells, including steroidogenic enzyme expression.Through gain-of-function experiments using miRNA mimics, we aimed to assess the impact of miR-6818-5p overexpression on granulosa cell proliferation and apoptosis, employing Cell Counting Kit-8 (CCK-8) assays and Annexin V/propidium iodide flow cytometry, respectively. To establish a direct mechanistic link, bioinformatic prediction was combined with experimental validation using dual-luciferase reporter assays to confirm targeting of candidate genes, complemented by Western blotting to assess corresponding protein expression changes. This multi-faceted methodology ensures a systematic investigation from phenotypic observation to molecular mechanism.

The primary objectives of this study are threefold. First, we aim to definitively characterize the expression pattern of miR-6818-5p in the peripheral blood of PCOS patients, establishing its potential as a circulating biomarker. Second, we seek to elucidate the functional role of miR-6818-5p in human ovarian granulosa cells, specifically evaluating its effects on proliferation and apoptosis. Third, we intend to identify and validate a direct target gene of miR-6818-5p that may mediate these cellular effects, thereby delineating a novel regulatory axis. By achieving these aims, this research endeavors to provide new insights into the molecular etiology of PCOS, moving beyond correlation to establish causality for a specific miRNA. The findings could pave the way for the development of miR-6818-5p as a diagnostic biomarker and lay the groundwork for future therapeutic strategies aimed at modulating miRNA activity to restore normal granulosa cell function in PCOS.

## 2. Materials and Methods

### 2.1 Clinical Samples and miRNA Extraction

Whole blood samples were collected from patients diagnosed with polycystic ovary syndrome (PCOS) according to the Rotterdam criteria (n = 5) and age-matched healthy controls (n = 5) at the Reproductive Medicine Center. All participants provided informed consent, and the study protocol was approved by the Institutional Ethics Committee. For mRNA quantification of miR-6818-5p expression efficiency, total cellular RNA was extracted using TRIzol reagent (Tiangen, cat. no. DP419) and reverse-transcribed with the PrimeScript RT Kit (Tiangen, cat. no. KR116-01), and miRNA concentrations were measured using a NanoDrop spectrophotometer.

### 2.2 Quantitative Real-Time PCR (qPCR)

Quantitative real-time PCR (qRT-PCR) was performed on a StepOnePlus Real-Time PCR System (Applied Biosystems). Amplification reactions were conducted using SYBR Premix Ex Taq (Tiangen, Cat. No. RK145) with specific primer pairs. U6 snRNA was selected as the internal reference gene for the relative quantification of miRNAs. Relative gene expression was calculated using the 2^−ΔΔCt^ method.

### 2.3 Cell Culture and Transfection

Human ovarian granulosa tumor (KGN) cells were cultured in Dulbecco’s Modified Eagle Medium/Nutrient Mixture F-12 (DMEM/F12) supplemented with 10% fetal bovine serum (FBS) and 1% penicillin-streptomycin at 37°C in a humidified atmosphere containing 5% CO2. For functional experiments, cells were seeded at appropriate densities and transfected with either miR-6818-5p mimics or mimic negative control (NC) using Lipofectamine 6000 (Beyotime, Cat. no. C0526) according to the manufacturer’s protocol and the transfection concentration was 20 nM. A lipofectamine-only group was included as an additional control for the assay.

### 2.4 Western Blotting Analysis

Total protein was extracted from KGN cells using RIPA lysis buffer(Beyotime, cat no. P0013C) supplemented with protease and phosphatase inhibitors. Protein concentrations were determined using the BCA Protein Assay Kit (Beyotime, cat no. P0012S). Equal amounts of protein were separated by SDS-PAGE on 10% gels and transferred to PVDF membranes. After blocking with 5% non-fat milk in TBST for 1 hour at room temperature, membranes were incubated overnight at 4°C with primary antibodies against HSD17B2 (1:1000; Sangon Biotech, cat no. D122504), and β-Tubulin (1:1000; Beyotime, cat no. AT819) as a loading control. Following three washes with TBST, membranes were incubated with HRP-conjugated secondary antibodies: Goat anti-mouse IgG (1:1000; Beyotime, cat no. A0216) and Goat anti-rabbit IgG (1:1000; Beyotime, cat no. A0208) for 1 hour at room temperature. Protein bands were visualized using enhanced chemiluminescence (ECL) substrate and quantified by densitometric analysis using ImageJ software. All Western blotting experiments were normalized to β-Tubulin expression

### 2.5 Dual-Luciferase Reporter Assay

The 3’-UTR sequence of HSD17B2 containing the predicted miR-6818-5p binding site was amplified by PCR and cloned into the pmirGLO Dual-Luciferase miRNA Target Expression Vector (Promega) downstream of the firefly luciferase gene to generate the wild-type (WT) construct. A mutant (Mut) construct was created by site-directed mutagenesis to disrupt the miR-6818-5p seed sequence complementarity. KGN cells were co-transfected with either the WT or Mut reporter plasmid together with miR-6818-5p mimics or mimic NC using Lipofectamine6000 (Beyotime, Cat. no. C0526). After 48 hours, firefly and Renilla luciferase activities were measured using the Dual-Luciferase Reporter Assay System (Promega, E1910) on a GloMax luminometer. Firefly luciferase activity was normalized to Renilla luciferase activity for each transfection group, and relative luciferase activity was calculated compared to the NC group.

### 2.6 Cell Proliferation Assay (CCK-8)

Cell proliferation was evaluated using the Cell Counting Kit-8 (CCK-8) assay(ZETA life, K009). KGN cells were seeded into 6-well plates and cultured to reach 80% confluence, followed by transfection with miR-6818-5p mimic or negative control mimic. Twenty-four hours after transfection, the cells were digested and reseeded into 96-well plates at a density of 5000 cells per well. At 6, 24, 48, 72 and 96 h after seeding, 10 μL of CCK-8 reagent was added to each well, followed by incubation of the plates at 37 °C for 2 h. Absorbance was measured at 450 nm using a microplate reader. All measurements were performed in six replicates per group per time point, and the experiment was independently repeated three times.

### 2.7 Flow Cytometry Apoptosis Analysis

Cell proliferation was evaluated using the Cell Counting Kit-8 (CCK-8) assay(ZETA life, K009). KGN cells were seeded into 6-well plates and cultured to reach 80% confluence, followed by transfection with miR-6818-5p mimic or negative control mimic.Twenty-four hours after transfection, the cells were digested and reseeded into 96-well plates at a density of 5000 cells per well. At 6, 24, 48, 72 and 96 h after seeding, 10 μL of CCK-8 reagent was added to each well, followed by incubation of the plates at 37 °C for 2 h. Absorbance was measured at 450 nm using a microplate reader. All measurements were performed in six replicates per group per time point, and the experiment was independently repeated three times.

### 2.8 Statistical Analysis

All experimental data are expressed as mean ± standard deviation (SD) from at least three independent experiments. Comparisons between two groups were performed using Student’s t-test. Statistical significance was defined as P < 0.05. All analyses were conducted using GraphPad Prism version 8.0 software.

## 3. Results

### 3.1 Elevated Expression of miR-6818-5p in PCOS Patients

As shown in Fig. 1, the expression level of miR-6818-5p in whole blood samples from PCOS patients was significantly elevated compared to that in healthy controls. qPCR quantification revealed that the average relative expression of miR-6818-5p in the PCOS group was approximately 1.59-fold higher than in the control group (P < 0.01). This marked upregulation of miR-6818-5p in the blood of PCOS patients suggests its potential involvement in the pathogenesis of the syndrome and positions it as a candidate biomarker for disease detection. The specificity of this alteration was further supported by normalization to U6 snRNA, which showed no significant variation between the two groups. These clinical data provide the foundation for subsequent cellular functional investigations to elucidate the mechanistic role of miR-6818-5p in ovarian granulosa cell biology.

**Figure 1.**
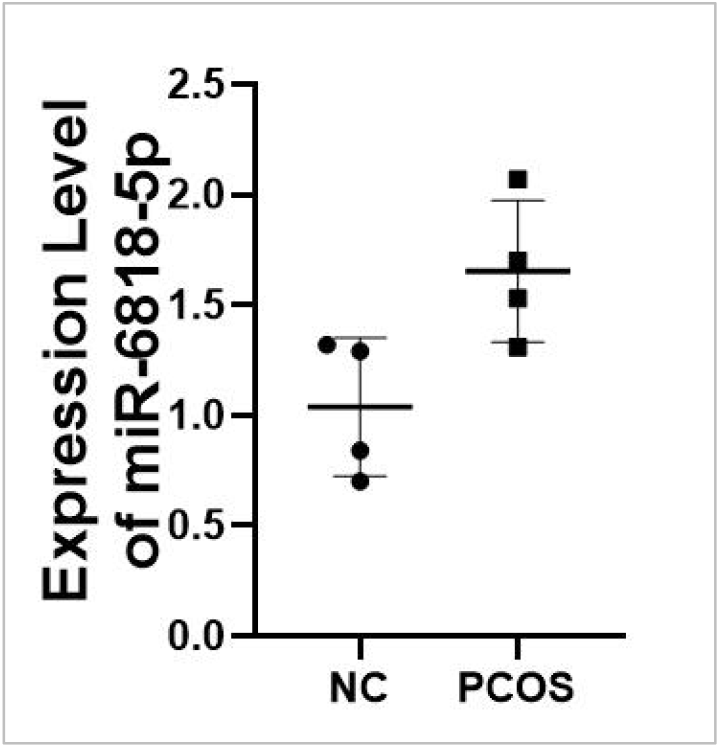
Analysis of miR-6818-5p expression in whole blood from PCOS patients by qPCR.qPCR quantification of exosomal miR-6818-5p levels in whole blood samples from PCOS patients compa red to healthy controls. Results demonstrate significant upregulation in the PCOS group (P<0.01), indicating its potential role in PCOS pathogenesis.

### 3.2 Successful Overexpression of miR-6818-5p in KGN Cells

As shown in Fig. 2, successful overexpression of miR-6818-5p was achieved in human ovarian granulosa KGN cells following transfection with miR-6818-5p mimics. qPCR analysis conducted 48 hours post-transfection demonstrated that the expression level of miR-6818-5p in the mimics group was approximately 180-fold higher than in the mimic NC group (P < 0.001). This substantial increase confirmed the efficiency of the transfection protocol and validated the experimental model for subsequent functional assays. The mimic NC group showed no significant deviation from baseline expression levels, indicating that observed effects in downstream experiments were attributable specifically to miR-6818-5p overexpression rather than non-specific transfection-related artifacts.

**Figure 2.**
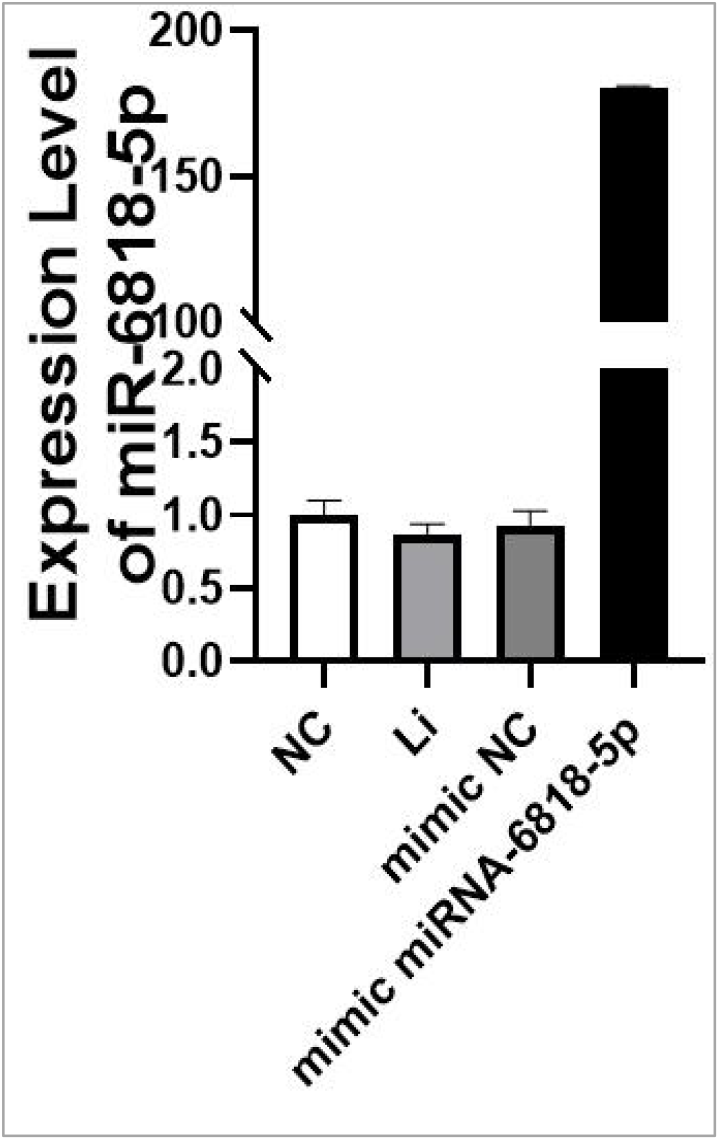
Validation of miR-6818-5p overexpression efficiency in KGN cells transfected with miRN A mimic-6818-5p.qPCR analysis confirming successful overexpression of miR-6818-5p in KGN cells following transfection with miRNA mimic-6818-5p versus mimic negative control (NC) group (P<0.001), establishing the model for functional assays.

### 3.3 HSD17B2 is a Direct Target of miR-6818-5p

As shown in Fig. 3, HSD17B2 was validated as a direct target gene of miR-6818-5p through both Western blotting and dual-luciferase reporter assays. Western blotting analysis (Fig. 3A) revealed that HSD17B2 protein expression in miR-6818-5p mimic-transfected KGN cells was reduced by approximately 6% compared to the mimic NC group, with β-Tubulin serving as the loading control to ensure equal protein loading across lanes. Densitometric quantification of the Western blot bands confirmed this significant downregulation, establishing that miR-6818-5p negatively regulates HSD17B2 at the protein level. To further verify the direct targeting relationship, a dual-luciferase reporter assay was employed (Fig. 3B). Co-transfection of the wild-type (WT) HSD17B2 3’-UTR reporter plasmid with miR-6818-5p mimics resulted in a approximately 28% reduction in relative luciferase activity compared to the mimic NC group. In contrast, the mutant (Mut) construct, featuring disrupted seed sequence complementarity, showed no significant change in luciferase activity upon co-transfection with miR-6818-5p mimics. This differential effect between WT and Mut constructs provides conclusive evidence that miR-6818-5p binds specifically to the predicted site within the HSD17B2 3’-UTR, thereby directly targeting HSD17B2 expression.

**Figure 3.**
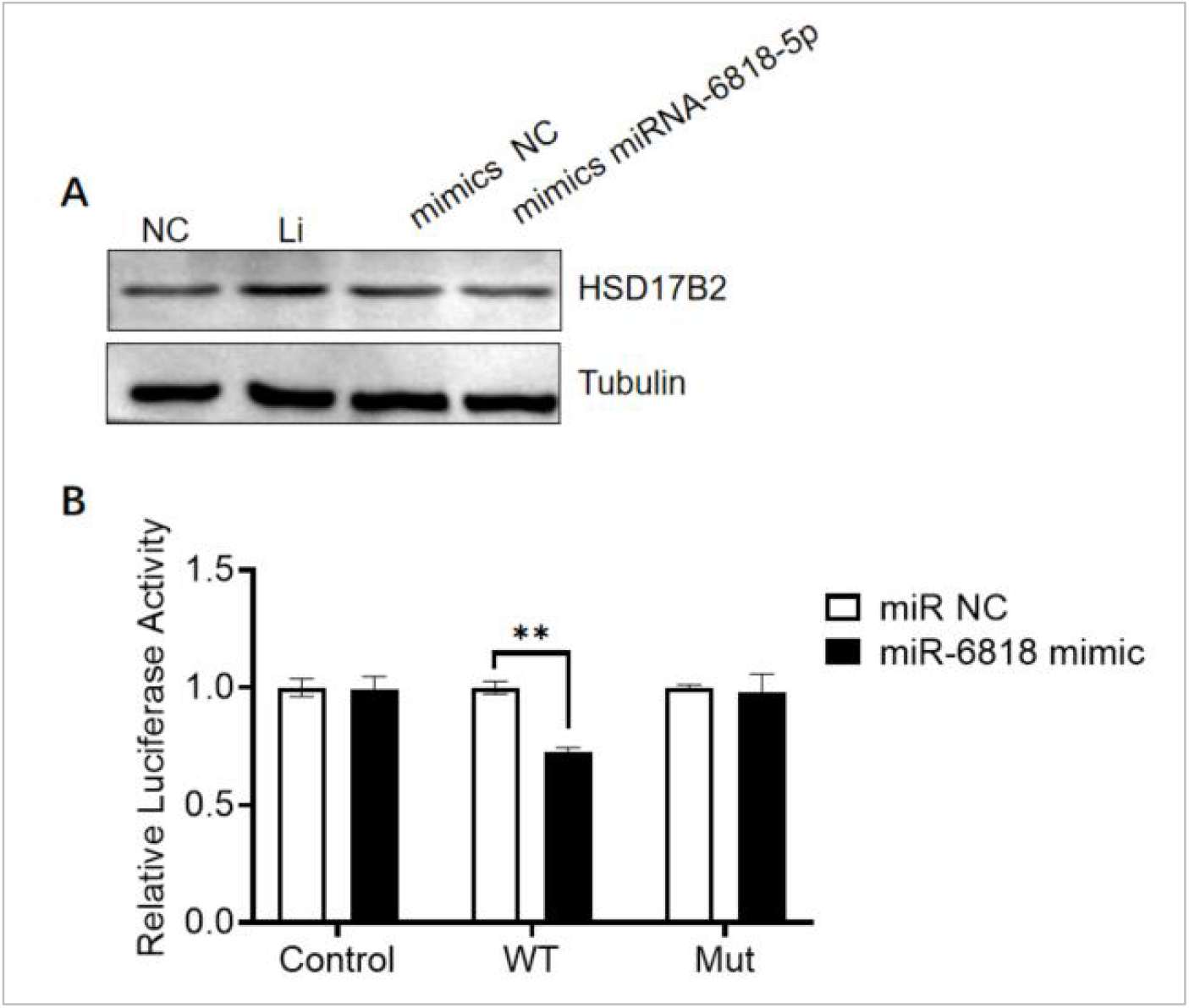
Target validation of HSD17B2 as a direct binding site for miR-6818-5p.(A) Western blot ting analysis showing reduced HSD17B2 protein expression in miR-6818-5p mimic-transfected KGN cells compared to NC. (B) Dual-luciferase reporter assay verifying direct targeting, with decreased luciferase activity in wild-type (WT) HSD17B2 constructs but not mutant (Mut) controls.

### 3.4 miR-6818-5p Overexpression Inhibits KGN Cell Proliferation

As shown in Fig. 4, CCK-8 proliferation assays revealed that miR-6818-5p overexpression significantly inhibited KGN cell proliferation. Cells were monitored at 6, 24, 48, 72 and 96 hours after cell seeding following transfection for 24 hours. While a modest decrease in cell viability was observed at 24-48 hours, the most pronounced effect occurred within 72-96 hours, where the proliferation rate of the miR-6818-5p mimics group was 76%-85% higher than that of the mimic NC group (P < 0.05). The absorbance values measured at 450 nm after 2-hour CCK-8 incubation consistently demonstrated reduced metabolic activity in the mimics group, correlating with decreased cell number.

**Figure 4.**
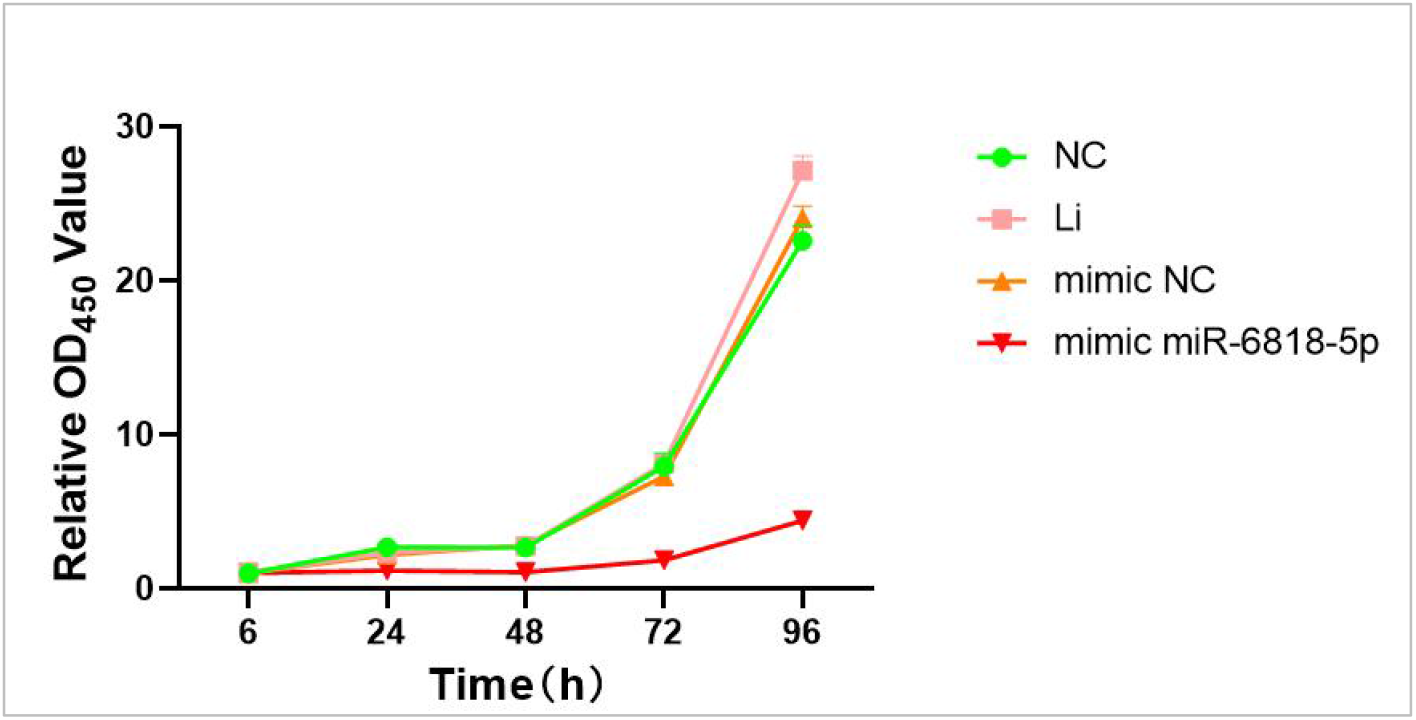
Proliferation assay of KGN cells overexpressing miR-6818-5p using CCK-8. Cell prolifera tion in each group was assessed via CCK-8 assay at 6, 24, 48, 72 and 96 h after cell seeding foll owing transfection. KGN cells (5,000 per well) overexpressing miR-6818-5p showed significantly red uced proliferation compared with the control group, detected at 450 nm after 2-hour incubation.

### 3.5 miR-6818-5p Overexpression Promotes KGN Cell Apoptosis

As shown in Fig. 5, flow cytometry analysis using Annexin V-FITC/PI dual staining demonstrated that miR-6818-5p overexpression significantly increased the apoptotic rate of KGN cells. At 48 hours post-transfection, the average apoptosis rate (sum of early apoptotic Q3 quadrant and late apoptotic Q2 quadrant populations) in the miR-6818-5p mimics group was approximately 11.83%, compared to approximately 9.67% in the mimic NC group, representing an increase of approximately 22% (P < 0.01). The lipofectamine-only control group showed an apoptotic rate comparable to the mimic NC group, confirming that the anti-apoptotic effect was specifically attributable to miR-6818-5p overexpression rather than to lipofectamine-mediated cellular stress. The pronounced reduction in apoptosis, particularly in the early apoptotic population (Q3 quadrant), is mechanistically consistent with the observed downregulation of Caspase-9-the initiator caspase of the intrinsic apoptotic pathway. These data complement the proliferation findings from Fig. 5, together demonstrating that miR-6818-5p exerts a dual regulatory effect on granulosa cell fate: suppressing proliferation while simultaneously promoting apoptosis through distinct yet interconnected signaling mechanisms.

**Figure 5.**
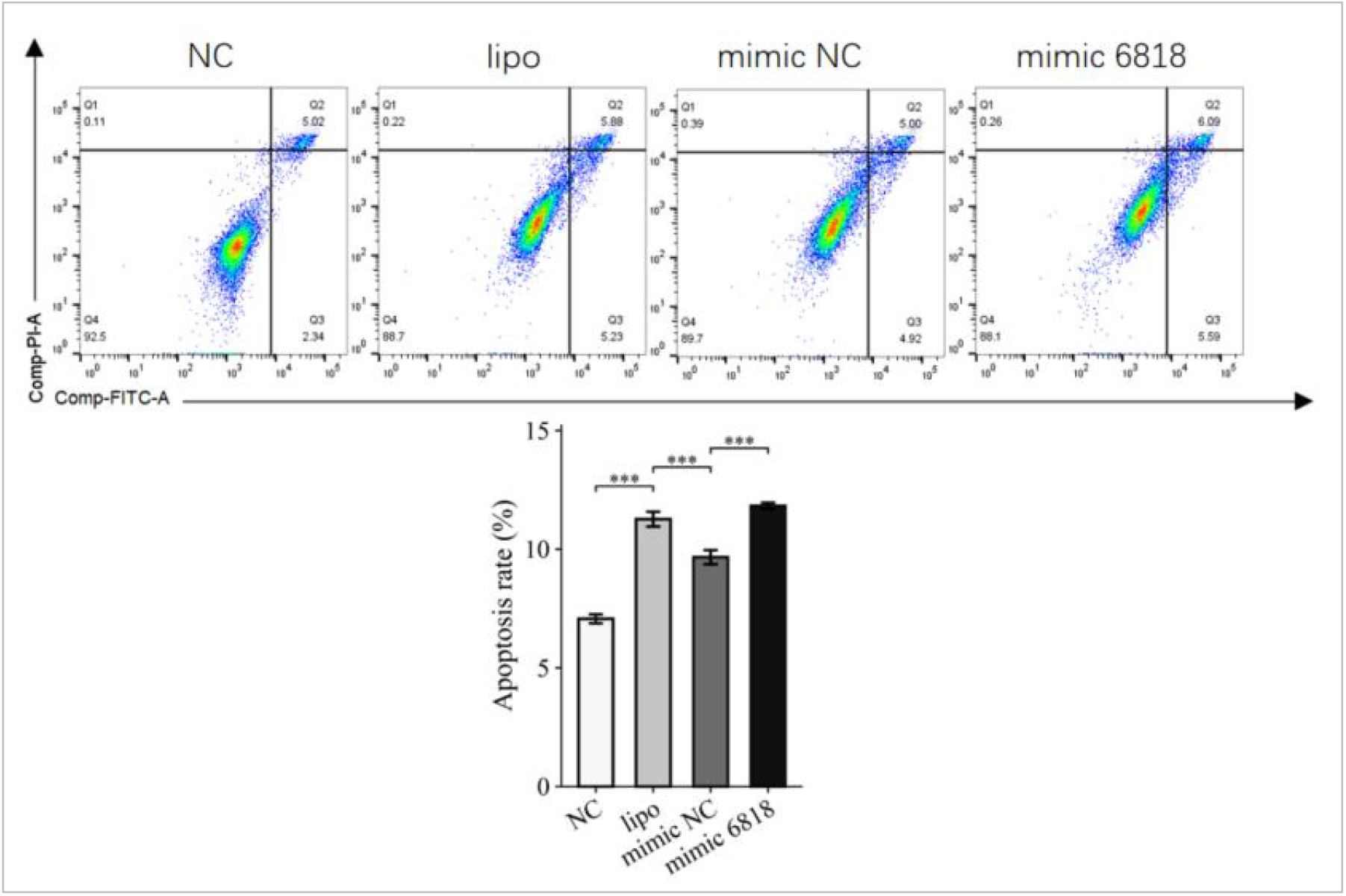
Flow cytometry analysis of apoptosis in KGN cells transfected with miR-6818-5p mimic. Transfection of miR-6818-5p mimic and flow cytometry to simulate apoptosis of KGN cells 48 hours after NC transfection, with NC group and lipofectamine group as controls. The simulation group showed a decrease in apoptosis rate, which was quantified by Annexin V/PI staining and Q2+Q 3 quadrant analysis.

## 4. Discussion

Polycystic ovary syndrome (PCOS) is a prevalent endocrine-metabolic disorder affecting women of reproductive age, characterized by hyperandrogenism, ovulatory dysfunction, and polycystic ovarian morphology, which profoundly impairs fertility and quality of life [1]. Although the precise pathogenesis of PCOS remains incompletely understood, dysfunction of ovarian granulosa cells (GCs) is recognized as a critical factor in aberrant folliculogenesis and anovulation. microRNAs (miRNAs), key post-transcriptional regulators, have been increasingly implicated in various diseases, modulating processes such as cell proliferation, apoptosis, and steroidogenesis. Despite this, the specific roles of miRNAs in PCOS-related GC dysfunction are poorly defined [8-9]. Notably, a high-throughput sequencing study identified miR-6818-5p as significantly upregulated in serum exosomes of PCOS patients, yet its functional significance and downstream targets in the context of PCOS pathology have remained unexplored [1].

This study was designed to elucidate the role of miR-6818-5p in PCOS by investigating its expression in patient blood and its functional impact on human granulosa cells. Our key findings reveal that miR-6818-5p is markedly elevated in the peripheral blood of PCOS patients, corroborating prior exosomal sequencing data [1], and that its overexpression in KGN granulosa cells directly suppresses the expression of HSD17B2, a critical enzyme in ovarian steroid metabolism. Functionally, this upregulation of miR-6818-5p potently inhibits cellular proliferation and induces apoptosis. These results identify a novel molecular axis—miR-6818-5p/HSD17B2—that links miRNA dysregulation to GC dysfunction, offering new insights into PCOS pathophysiology. The subsequent discussion will delve into the potential clinical significance of circulating miR-6818-5p, the mechanistic implications of its targeting of HSD17B2 within the steroidogenic pathway, and the broader cellular consequences of its aberrant expression.

The present study identified a significant upregulation of circulating miR-6818-5p in the peripheral blood of PCOS patients compared to healthy controls, a finding that aligns with a prior serum exosomal miRNA sequencing study which also reported miR-6818-5p as a differentially expressed candidate in PCOS [1]. This concordance across different sample types—whole blood and serum exosomes—strengthens the evidence for this miRNA’s involvement in the disease. The observed elevation is unlikely to be a mere bystander effect; it may reflect a systemic pathological state or a local ovarian response. Given that granulosa cells are the primary functional unit within the follicle, the increase in circulating miR-6818-5p could originate, at least in part, from dysfunctional or apoptotic granulosa cells themselves, mirroring the cellular stress we observed in vitro. Furthermore, other reported dysregulated miRNAs in PCOS, such as miR-21 which targets TLR8 [10] and miR-135b which targets LATS2 [11], have been linked to inflammation and Hippo signaling, respectively. In contrast, our result suggests a unique association of miR-6818-5p with the downstream steroidogenic pathway, potentially adding a complementary layer to the complex miRNA regulatory network in PCOS. While this finding establishes a clinical correlate, the functional significance of this 1.5-fold change in circulation, its correlation with specific patient phenotypes such as hyperandrogenism or insulin resistance, and its cell-type specific origin warrant further investigation in larger, phenotypically well-characterized cohorts. This initial discovery firmly positions miR-6818-5p as a candidate for further exploration in the molecular pathology of PCOS.

The post-transcriptional suppression of HSD17B2 by miR-6818-5p is therefore predicted to dysregulate the intraovarian steroid environment, potentially leading to a localized accumulation of androgens. This mechanistic link is particularly compelling given that hyperandrogenism is a cornerstone of PCOS pathology. Mechanistically, our dual-luciferase reporter and Western blotting experiments confirmed HSD17B2 as a direct and functionally relevant target of miR-6818-5p.HSD17B2 encodes 17β-hydroxysteroid dehydrogenase type 2, an enzyme that catalyzes the conversion of the potent androgen testosterone to the weaker androgen androstenedione, and of estradiol to estrone, thereby playing a crucial role in attenuating local androgen and estrogen activity within the follicle[12-14].

While our findings focus on this single target, miRNAs typically regulate hundreds of transcripts. It is plausible that miR-6818-5p also modulates other mRNAs relevant to PCOS, such as those involved in insulin signaling or cell cycle regulation, consistent with the multi-target regulatory patterns described for other miRNAs [11]. The miR-6818-5p/HSD17B2 axis we identified thus provides a direct molecular bridge from a dysregulated miRNA to a critical enzyme in steroid hormone metabolism, offering a new mechanistic perspective on how post-transcriptional control may contribute to the classic endocrine imbalance of PCOS. This specific regulatory relationship highlights a previously unappreciated node in the molecular network governing ovarian steroidogenesis.

The biological consequence of this molecular event was demonstrated by our functional assays, which showed that miR-6818-5p overexpression significantly inhibited proliferation and promoted apoptosis in KGN granulosa cells. This phenotype is consistent with the concept that suppressed granulosa cell proliferation and increased atresia are key features of follicular dysfunction in PCOS. In our context, the reduction of HSD17B2 protein is a plausible upstream event. Diminished HSD17B2 activity would disrupt local steroid conversion; elevated intrafollicular androgens, for instance, are known to trigger granulosa cell apoptosis and follicular atresia. This proposition is supported by studies showing that alterations in signaling pathways, such as the PI3K/Akt/mTOR pathway, mediate apoptosis in rat granulosa cells [15] and that other regulatory molecules like DLGAP5 anti-apoptosis effects are altered in PCOS [16]. In our system, the pro-apoptotic and anti-proliferative effects of miR-6818-5p are therefore likely mediated through the downstream consequences of HSD17B2 suppression, rather than a direct interaction with a cell cycle regulator. It remains to be determined whether the observed reduction in proliferation is a direct result of cell cycle arrest or a secondary consequence of the induction of apoptosis, a question that could be resolved by examining cell cycle phase distribution. These results establish a cellular function for miR-6818-5p and mechanistically connect it to a specific pathological outcome, providing a plausible explanation for the impaired follicle development observed in PCOS.

Finally, the functional role of miR-6818-5p, as defined by our gain-of-function experiments, establishes it as a novel negative regulator of granulosa cell function. Its elevated expression in PCOS patients positions it not merely as a biomarker but as a potential contributor to the disease pathology by actively suppressing HSD17B2 and subsequently driving apoptosis. This stands in contrast to other reported miRNA functions in PCOS; for example, miR-135b overexpression was shown to promote granulosa cell proliferation by inhibiting the Hippo pathway [11], while miR-21’s upregulation was linked to increased inflammation [10]. Our results identify miR-6818-5p as belonging to the category of miRNAs that are detrimental to granulosa cell survival. The strength of this functional conclusion would be substantially augmented by loss-of-function experiments using specific inhibitors (antagomirs) in patient-derived granulosa cells. If such an intervention could rescue the proliferative capacity and reduce the apoptotic rate, it would provide strong evidence for the pathophysiological necessity of miR-6818-5p upregulation. Furthermore, discerning whether this elevated expression represents an exaggerated version of a normal physiological role during folliculogenesis or an entirely aberrant pathological event is crucial. These findings firmly establish the functional importance of this miRNA and underscore its potential as a therapeutic target, though the effectiveness and specificity of such an approach require extensive preclinical validation.

Several limitations of this study warrant consideration. First, the clinical validation cohort remains small (n=5 per group), limiting statistical power and generalizability—the observed 1.5-fold upregulation of miR-6818-5p in PCOS patients requires independent replication in larger, multi-center cohorts to confirm its diagnostic utility and assess potential correlations with specific phenotypes (e.g., hyperandrogenism or insulin resistance). Second, the mechanistic experiments relied exclusively on the KGN granulosa cell line, which may not fully recapitulate the physiology of primary human granulosa cells or the heterogeneous ovarian environment in PCOS patients; the absence of in vivo validation using a PCOS animal model (e.g., DHEA-induced rats) precludes definitive conclusions about the physiological relevance of the miR-6818-5p/HSD17B2 axis. Additionally, the study lacked loss-of-function experiments (e.g., miR-6818-5p inhibition), leaving unanswered whether reversing its upregulation could restore HSD17B2 expression and normalize granulosa cell function—a critical step for establishing causal necessity.

In summary, this study identifies, for the first time, elevated circulating miR-6818-5p in PCOS patients, and demonstrates its direct targeting of HSD17B2, leading to suppressed proliferation and enhanced apoptosis in KGN cells. These findings establish a novel miR-6818-5p/HSD17B2 axis that may contribute to PCOS-associated follicular dysfunction, offering potential diagnostic biomarkers (circulating miR-6818-5p) and therapeutic targets (e.g., antagomirs). Future work should prioritize large-scale clinical validation, in vivo confirmation using animal models, and functional loss-of-rescue experiments to translate these molecular insights into clinical applications for PCOS management.

